# Comprehensive elucidation of glutathione import in *Escherichia coli*

**DOI:** 10.1101/2024.07.15.603537

**Authors:** Lisa R. Knoke, Maik Muskietorz, Lena Kühn, Lars I. Leichert

## Abstract

Glutathione is the major thiol-based antioxidant in a wide variety of biological systems, ranging from bacteria to eukaryotes. As a redox couple, consisting of reduced glutathione (GSH) and oxidized glutathione disulfide (GSSG), it is crucial for the maintenance of the cellular redox balance. Glutathione transport out of and into cellular compartments and the extracellular space is a determinant of the thiol-disulfide redox state of the organelles and bodily fluids in question, but is currently not well understood. Here we use the genetically-encoded, glutathione-measuring redox probe Grx1-roGFP2 to comprehensively elucidate the import of extracellular glutathione into the cytoplasm of the model organism *Escherichia coli*. The elimination of only two ATP-Binding Cassette (ABC) transporter systems, Gsi and Opp, completely abrogates glutathione import into *E. coli*’s cytoplasm, both in its reduced and oxidized form. The lack of only one of them, Gsi, completely prevents import of oxidized glutathione (GSSG), while the lack of the other, Opp, substantially retards the uptake of reduced glutathione (GSH).

## Introduction

The tripeptide glutathione (GSH), noted for its reductive properties, is one of the most abundant antioxidants in various organisms ranging from bacteria to eukaryotes (Fahey et al., 1978; Smirnova and Oktyabrsky, 2005; Zechmann et al., 2011). In the cytosol of *E. coli*, GSH is present in the millimolar range with concentrations of 5-10 mM and the [GSH]/[GSSG] ratio has been determined to be 50:1 to 200:1. GSH is actively maintained in this reduced state by the glutathione reductase (Gor), which uses NADPH as reductant (Åslund et al., 1999; Greer and Perham, 1986; Masip et al., 2006; Meister, 1988). Protein oxidation in the cytosol of cells often results in a loss of function. Thus GSH, in concert with the Glutaredoxins (Grx), is important for protein reduction in Gram-negative bacteria such as *E. coli* (another important pathway being the thioredoxin pathway). Grxs depend on reduced GSH for reduction of their active site. GSH, when oxidized, dimerizes to GSSG, the glutathione disulfide (Aslund et al., 1994; Carmel-Harel and Storz, 2000). In *E. coli,* the tripeptide GSH is synthesized in the cytoplasm in an ATP-dependent two step mechanism. For that, the γ-glutamate-cysteine ligase (GshA) condenses the carboxyl group at the γ-position of glutamate with the amino group of cysteine, creating the dipeptide γ-glutamyl-cysteine. In a second step, glutathione synthase (GshB) adds a glycine to the C-terminus of this dipeptide, resulting in γ-glutamyl-cysteinyl-glycine (glutathione) (Apontoweil and Berends, 1975; Fuchs and Warner, 1975; Richman and Meister, 1975).

*E. coli* cells actively cycle GSH into the culture medium and back into their cytosol (Smirnova et al., 2012). Glutathione, like other small molecules, is presumably transported across the outer membrane through porins, but transport through the inner cell membrane requires specific ABC-transporters (Pittman et al., 2005; Smirnova et al., 2012; Suzuki et al., 2005). Previous studies showed that the ABC transporter Gsi composed of GsiA-D confers GSH import across the inner membrane into the cytosol (Suzuki et al., 2005; Wang et al., 2018). It was recently shown that loss of this glutathione transport system in the Gram-negative *Cronobacter sakazakii* led to a reduced desiccation tolerance and decreased intracellular glutathione (Wang et al., 2024). ABC transporters are important for import and export of a variety of different substrates, such as amino acids, sugars and peptides (Berntsson et al., 2010). They consist of a periplasmic binding protein that captures the transported substrates in the periplasm and targets them to the two inner membrane-spanning permease domains. The cytosolic ATPase domain(s) provide energy for the transport through ATP hydrolysis (Davidson and Chen, 2004; Higgins and Linton, 2004).

Here, we explore the import of glutathione in mutants lacking endogenous glutathione and components of known peptide transporters. In these mutants we express a genetically encoded redox-sensitive probe fused to glutaredoxin that allows us to track changes in the cytosolic glutathione state in real-time. We provide evidence that glutathione import *in vivo* is different from our current model. Our data indicates that Gsi is the major transporter of oxidized, but not reduced glutathione. The majority of reduced glutathione is instead transported through the peptide transporter Opp. Gsi and Opp are the only transporters for exogenous glutathione in *E. coli*.

## Materials and methods

### Strains, Plasmids and growth conditions

The strains used in this study are listed in Table 1 and plasmids in Table 2. All *E. coli* strains used originated or were derived from the KEIO collection (Baba et al., 2006), and were routinely cultivated at 37 °C in Luria-Bertani (LB) medium, supplemented with antibiotics when required for plasmid maintenance and marker selection (ampicillin 200 μg/mL or kanamycin 100 μg/mL), if not stated differently.

**Table 1.**
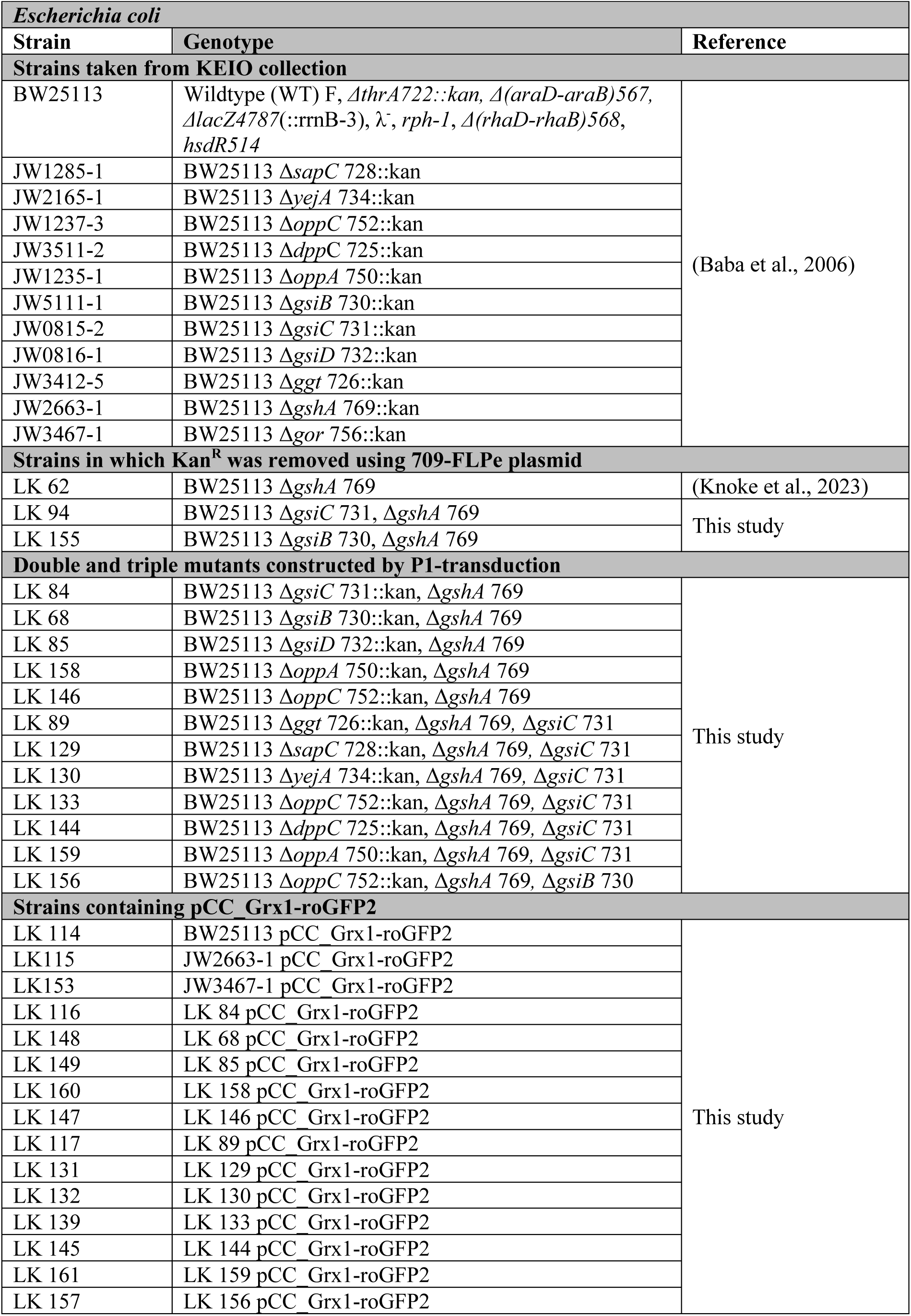
Bacterial strains used in this study.

**Table 2.**
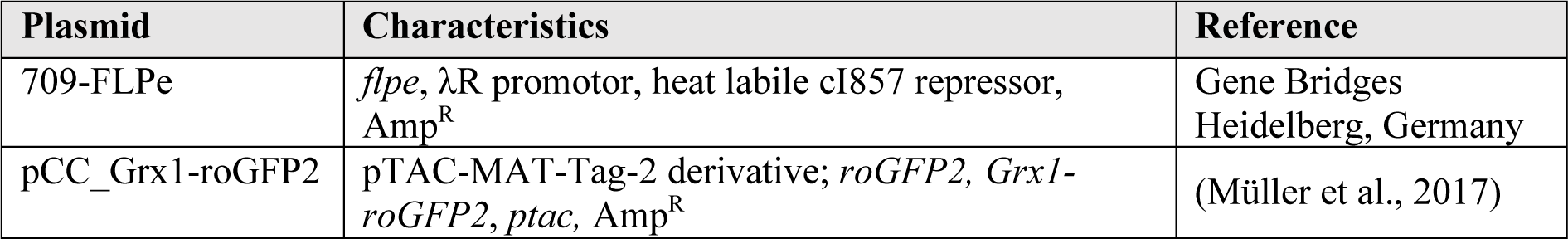
Recombinant plasmids used in this study.

| Plasmid | Characteristics | Reference |
| --- | --- | --- |
| 709-FLPe | *flpe*, λR promotor, heat labile cI857 repressor, Amp<sup>R</sup> | Gene Bridges Heidelberg, Germany |
| pCC_Grx1-roGFP2 | pTAC-MAT-Tag-2 derivative; *roGFP2*, *Grx1-roGFP2*, *ptac*, Amp<sup>R</sup> | (Müller et al., 2017) |

For expression of Grx1-roGFP2, mutant strains (Table 1) carrying the pCC_Grx1-roGFP2 plasmid (Table 2) were grown at 37 °C in MOPS minimal medium (Neidhardt, Teknova, Hollister, CA, USA) to an optical density (OD) of ∼0.5. Then, Grx1-roGFP2 expression was induced with 0.2 mM IPTG (Isopropyl ß-D-1-thiogalactopyranoside) and cells were further incubated for 16 h at 20 °C.

### Construction of E. coli gshA double and triple deletion strains using P1 transduction

For the construction of *E. coli gshA* double mutants, the *gshA* KEIO mutant without kanamycin cassette generated previously (Knoke et al., 2023) was used as acceptor for P1 transduction as described previously (Thomason et al., 2007). The KEIO *gsiB* (JW5113-1), *gsiC* (JW0815-2), *gsiD* (JW0816-1), *oppA* (JW1235-1) and *oppC* (JW1237-3) mutants were used as donors for P1 transduction. Subsequently, correct insertion of the mutant gene fragments into the marker free *gshA* mutant were checked using appropriate forward and reverse primers of the respective genes in combination with the *k1* and *k2* KEIO primers listed in Table 3 using colony PCR. For the generation of triple mutants, the kanamycin cassette was removed from the *gshA, gsiC* or *gshA, gsiB* double deletion mutant using the plasmid 709-FLPe as indicated by the supplier (Gene Bridges, Heidelberg, Germany).

**Table 3.**
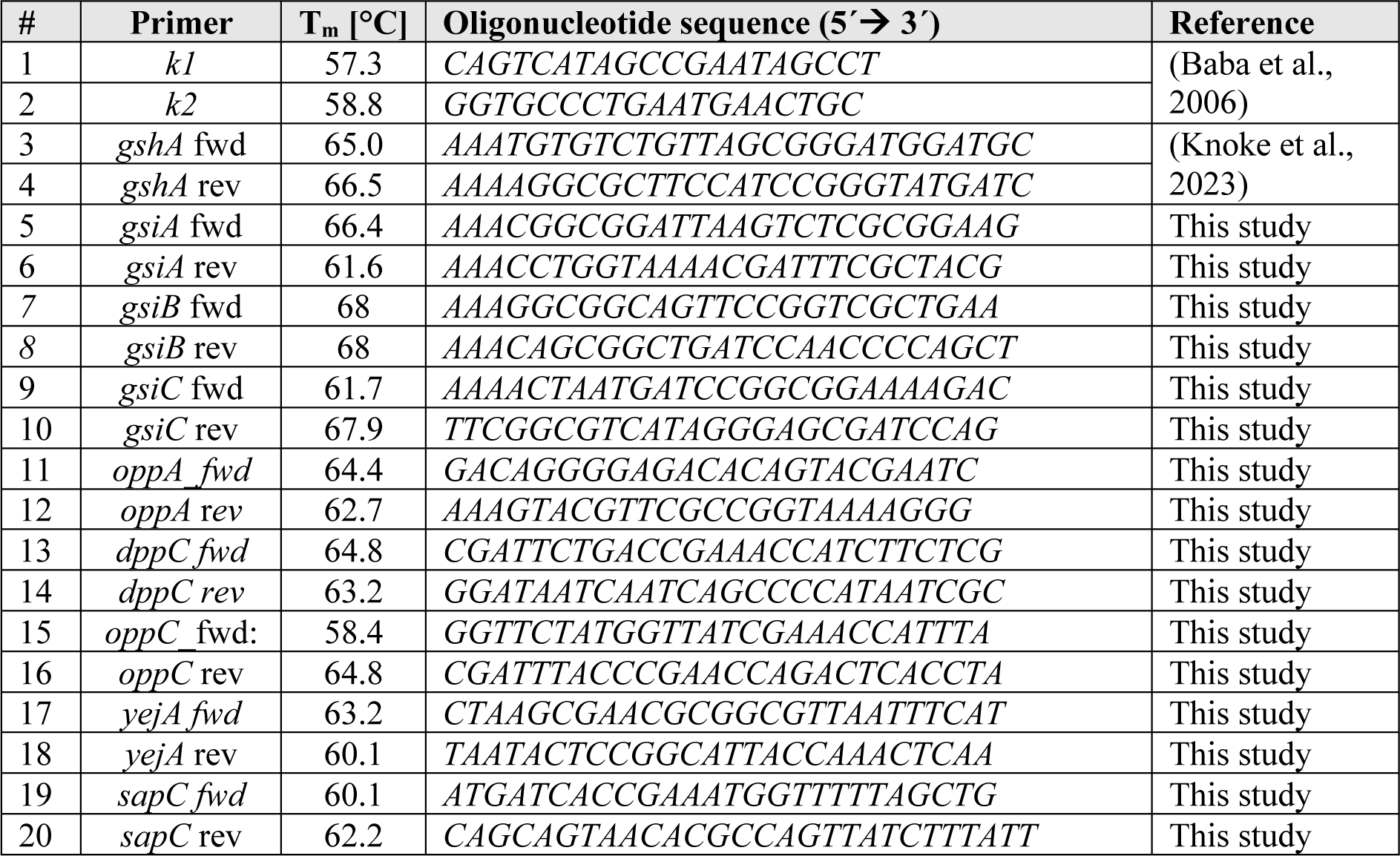
Oligonucleotides used in this study.

The removal of the kanamycin cassette was verified by colony PCR using *k1, k2* and gene specific primers (Table 3). The *oppA, oppC, ggt* (JW2919-1)*, sapC* (JW1285-1)*, yejA* (JW2165-1) or *dppC* (JW3511-2) mutants from the KEIO collection were used as donors for P1 transduction. Again, correct insertion of the KEIO fragments in the marker free double mutants was verified using colony PCR with appropriate primers (Table 3). All mutants were transformed with the pCC_Grx1-roGFP2 plasmid by heat shock and selected for ampicillin and kanamycin resistance.

### Selection of Gsi homologs in the search for additional glutathione transporters

The protein sequence of GsiB from *Escherichia coli* K12 (uniprot accession P75797) was used as a template for a BLASTp (Altschul et al., 1990) query against the *Escherichia coli* K12 substr. MG1655 reference genome found in the EcoCyc database (Karp et al., 2023). Expectation Value Threshold was set to 10, Matrix used was BLOSUM62 with a “Gap existence cost” of 11, a “Per residue gap cost” of 1 and a “Lambda ratio” of 0.85. No “Other advanced options” were chosen and the “Filter query sequence for low complexity regions” was set to default. Reported “Positives” in the results were assumed as similarity in %. Based on these results, the Dpp-, Opp-, Sap-and Yej-ABC transporters were selected for further testing.

### Grx1-roGFP2-based measurements in E. coli

The Grx1-roGFP2 oxidation state was determined as described before, with minor changes (Degrossoli et al., 2018; Knoke et al., 2023; Xie et al., 2020). Briefly, for determination of the Grx1-roGFP2 oxidation state in different *E. coli* mutant cells (Table 1), cells were transformed with Grx1-roGFP2 coding plasmids, and after expression of the probes for 16 h at 20 °C, as mentioned above, cells were harvested, washed twice in HEPES (40 mM, pH 7.4) and adjusted to an OD_600_ of 2.0 in HEPES. Then 50 µL of the cells were transferred to the wells of a black, clear-bottom 96-well plate (Nunc, Rochester, NY). Two-fold concentrated stock solutions containing either 2 mM Aldrithriol-2 (Sigma-Aldrich, Darmstadt, Germany, CAS-2127-03-9, AT-2) (oxidation control), 10 mM Dithiothreitol (Sigma-Aldrich, Darmstadt, Germany, CAS-3483-12-3, DTT) (reduction control), 5 mM reduced glutathione (GSH, Sigma-Aldrich, CAS-70-18-8, pH 7.4 in 1 M HEPES), 5 mM oxidized Glutathione (GSSG, Sigma-Aldrich, CAS-27025-41-8, pH 7.4 in 1 M HEPES) or HEPES were prepared freshly. For analysis of EDTA and Gor/NADPH influence on GSH import in cells lacking GshA and OppC, 10 mM EDTA or 4 µM Gor/500 µM NADPH with or without 5 mM GSH were used. DMSO, the solvent of AT-2 was added to all stocks, accounting for changes in oxidation based on DMSO. Then, 50 µL of the two-fold stocks were added to the cells immediately before fluorescence intensities were recorded in a CLARIOStar Plus (BMG Labtech, Germany) microplate reader. Changes of Grx1-roGFP2 were measured at excitation wavelengths 405 and 488 nm (bandwidth: 5 nm) and emission at 525 nm (bandwidth: 5 nm) at 25 °C.

Subsequently, the probe’s oxidation state was calculated as described before (Knoke et al., 2023; Xie et al., 2020) from the ratios of the fluorescence excitation intensities (405/488 nm). All values were normalized to fully oxidized (AT-2-treated) and fully reduced (DTT-treated) cells with the following equation [1]:

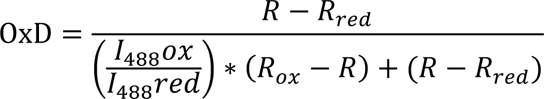

with *R_ox_* being the 405/488 nm ratio of oxidized (AT2-treated) and *R_red_* of reduced (DTT-treated) cells respectively. *I*_488_*ox* and *I*_488_*red* are the fluorescence intensities of Grx1-roGFP2 at 488 nm under oxidizing or reducing conditions. *R* is the measured 405/488 nm ratio.

Data was processed using Microsoft Excel software and graphs were generated using GraphPad Prism.

## Results

### Oxidized or reduced glutathione reduces Grx1-roGFP2 in cells lacking glutathione biosynthesis, but not in glutathione reductase-deficient cells

The genetically encoded redox probe roGFP2 contains two cysteines that, when oxidized, form a disulfide, changing the excitation properties of the probe. This allows for a ratiometric and concentration-independent determination of the probe’s dithiol disulfide state by fluorescence measurements. Unfused roGFP2 can react with cellular antioxidant systems, such as thioredoxin and glutaredoxin (Grx). Particularly the latter is an efficient thiol-disulfide oxidoreductase for roGFP2. However, the velocity of this reaction is limited by the number and vicinity of endogenous Grx (see Figure 1A for a schematic representation). (Dooley et al., 2004; Lukyanov and Belousov, 2014). Fusion of human glutaredoxin 1 (Grx1) to roGFP2 overcomes this kinetic barrier and brings it into equilibrium with the circumjacent glutathione pool, essentially making Grx1-roGFP2 a highly sensitive probe for the GSH/GSSG redox state of the cell (Gutscher et al., 2008; Meyer and Dick, 2010) (see Figure 1B for a schematic representation).

**Figure 1.**
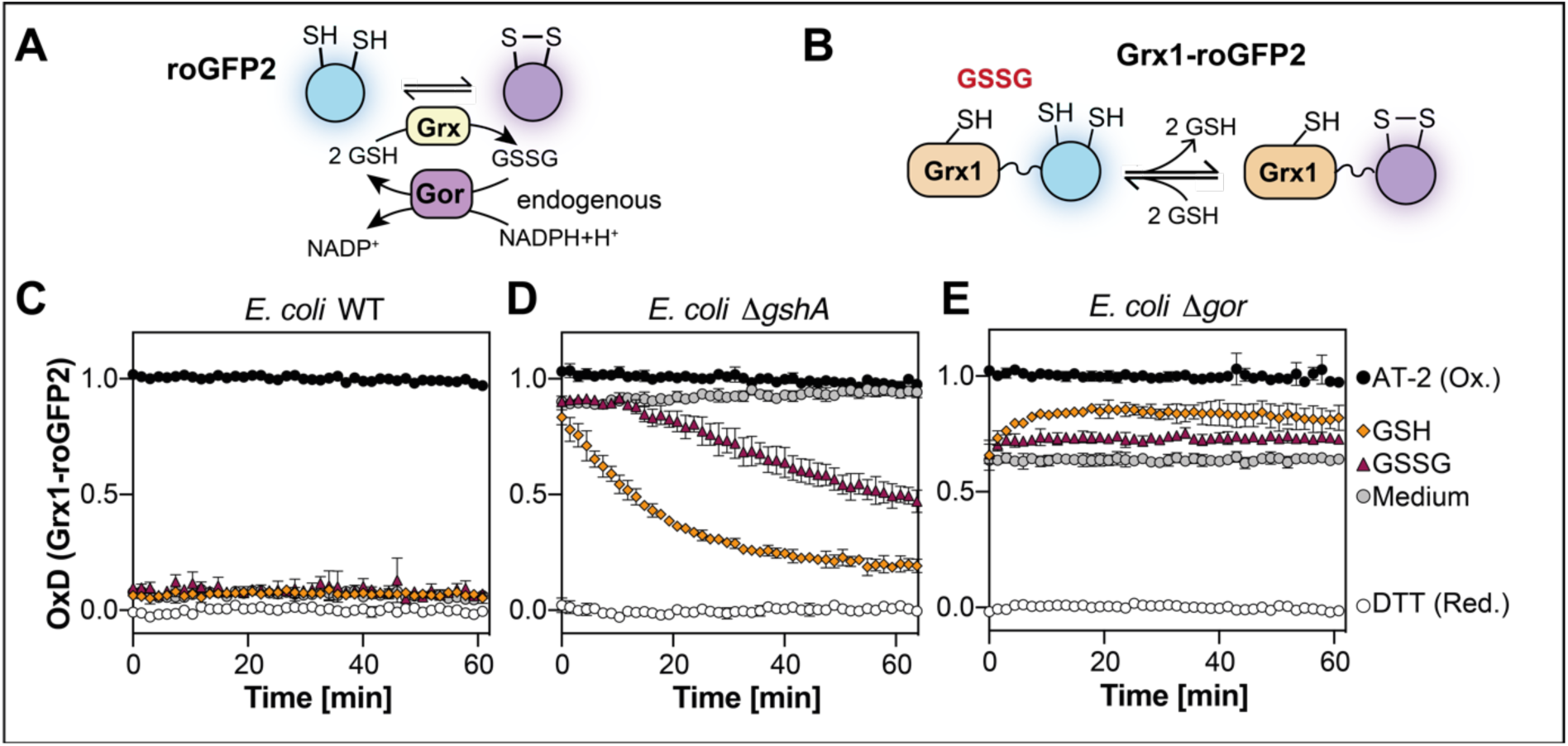
External glutathione reduces Grx1-roGFP2 in cells lacking endogenous glutathione biosynthesis but not in glutathione reductase deficient cells. Schematic overview of roGFP2’s **(A)** and Grx1-roGFP2’s **(B)** response to glutathione. Oxidation state of Grx1-roGFP2 expressed in *E. coli* WT **(C)**, Δ*gshA* **(D)** or Δ*gor* **(E)** during steady state conditions (medium) and in response to exogenous GSH and GSSG. Cells expressing the probes were cultivated for 16 h at 20 °C in MOPS minimal medium without glutathione, harvested and washed. Probe oxidation was then measured in 40 mM HEPES pH 7.4 supplemented without (medium) or with 2.5 mM GSH or GSSG. Grx1-roGFP2 oxidation was recorded at 525 nm emission and 405 nm and 488 nm excitation. Oxidized (AT-2) and reduced (DTT) cells were used as controls for calculation of the probe’s oxidation degree (OxD). Represented values are the mean of three independent experiments. All experiments were performed as technical triplicates.

We, thus, used Grx1-roGFP2 as a tool to monitor the GSH/GSSG redox state of *E. coli*’s cytoplasm. In the cytoplasm, Grx1-roGFP2 was virtually completely reduced, reflecting the well-known fact that cells typically have a highly reduced cytosolic glutathione pool. Addition of exogenous GSH or GSSG did not cause any measurable changes in the cellular GSH/GSSG homeostasis (Figure 1C).

But when we expressed the probe in the cytosol of cells lacking the first enzyme for glutathione biosynthesis (Δ*gshA*), Grx1-roGFP2 was virtually completely oxidized. This suggested to us that *E. coli* cannot maintain reduction of Grx1-roGFP2 in the absence of this antioxidant tripeptide. However, external addition of reduced glutathione (GSH) resulted in reduction of the probe. This shows that *E. coli* is able to both take up and utilize glutathione from the surrounding media. Somewhat counterintuitively, the addition of oxidized glutathione (GSSG) also resulted in probe reduction. *In vitro*, GSSG addition leads to oxidation of Grx1-roGFP2 (Müller et al., 2017). This observation suggests that *E. coli* can also take up oxidized glutathione from the surrounding media and then put it to use as an antioxidant by converting it to reduced glutathione. The observed delay of probe reduction by GSSG could be due to the time endogenous glutathione reductase needs to reduce the imported GSSG (Figure 1D). And indeed, in cells lacking the glutathione reductase (Gor) (Δ*gor*), Grx1-roGFP2 was around 60 % oxidized and neither addition of GSH, nor GSSG resulted in reduction of the probe (Figure 1E).

### The GsiA-D ABC transporter is essential for import of oxidized, but not reduced glutathione

In *E. coli* one transporter for import of the tripeptide glutathione into the cytoplasm, namely Gsi, consisting of GsiA-D, has been identified (see Figure 2A for a schematic overview) (Smirnova et al., 2012; Suzuki et al., 2005; Wang et al., 2018). We wondered if this transporter was the only factor that can convey the uptake of extracellular glutathione. To address this, we generated double deletion strains lacking both Δ*gshA* and either the gene coding for the periplasmic binding protein GsiB, or one of the two permease domains coding genes *gsiC* or *gsiD*. We then expressed Grx1-roGFP2 in the resulting double deletion strains and determined reduction of the probe after the addition of GSH. Deletion of any component of the GsiA-D transporter did not prevent Grx1-roGFP2 reduction by GSH (Figure 2B-D). This suggests that Gsi is not the only transporter for GSH. However, GSSG supplementation did no longer lead to reduction of Grx1-roGFP2 in these strains, independent on whether the periplasmic binding protein GsiB or the permease domains GsiC or GsiD were missing, suggesting that Gsi is the exclusive transporter for oxidized glutathione in *E. coli* (Figure 2B-D).

**Figure 2.**
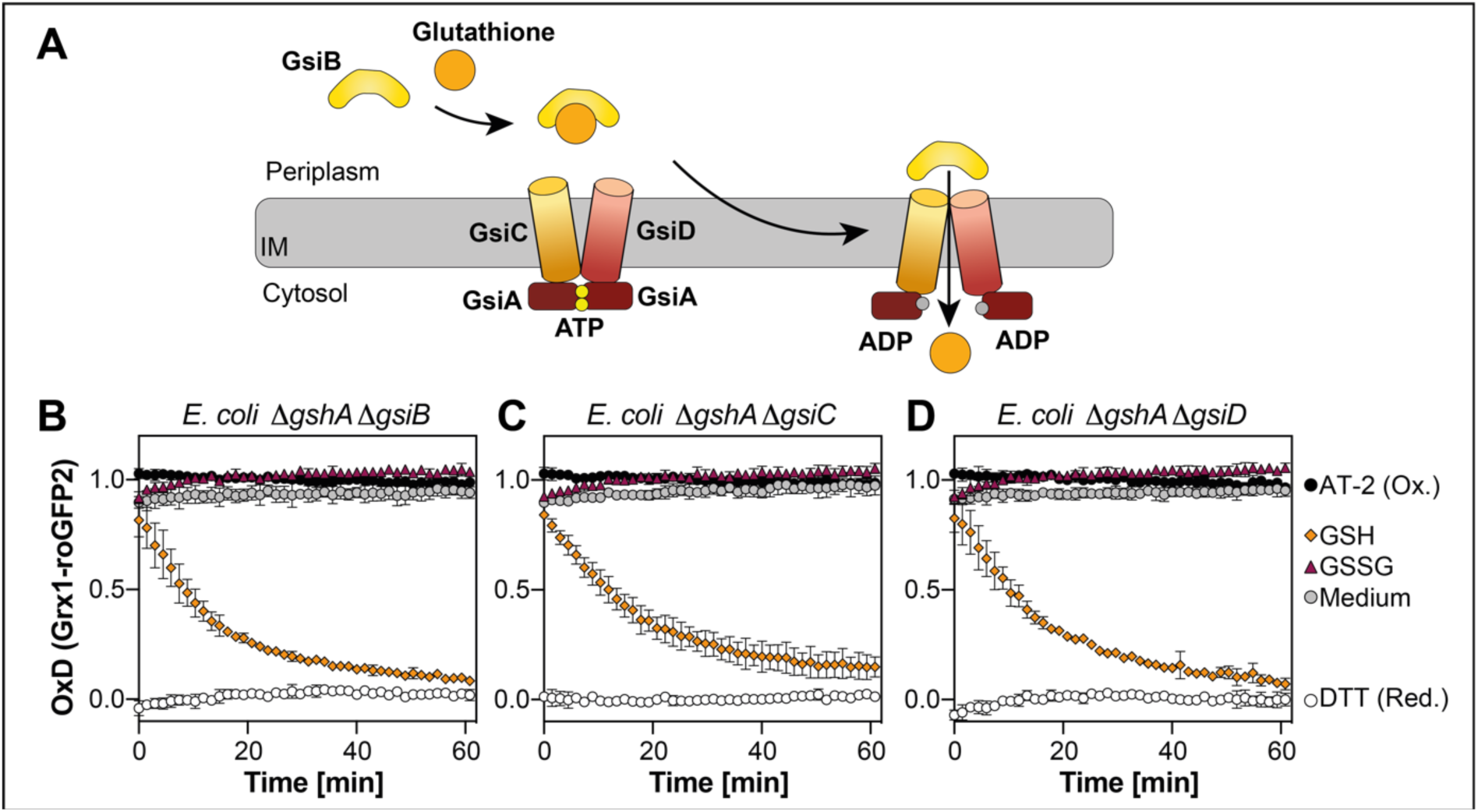
GsiA-D is the sole transporter for oxidized glutathione in *E. coli*. **(A)** Schematic overview of glutathione import by the Gsi ABC transporter. Grx1-roGFP2 oxidation state expressed in *E. coli* Δ*gshA* Δ*gsiB* **(B)**, Δ*gshA* Δ*gsiC* **(C)** or Δ*gshA* Δ*gsiD* **(D)**. Grx1-roGFP2 probe was expressed and its oxidation was recorded as described in Figure 1. Oxidized (AT-2-) and reduced (DTT-treated) cells were used for calculation of the probe’s oxidation degree (OxD). Plotted values represent the mean of three individual experiments, error bars represent the standard deviation. All experiments were performed as technical triplicates. *Glutathione hydrolysis by GGT is not the reason for the reduction of Grx1-roGFP2 in Gsi-deficient cells*

It has been suggested that most of the glutathione taken up from the environment into the cytoplasm is not transported as a tripeptide, but degraded by the inner membrane bound glutathione transpeptidase Ggt. After hydrolysis, the resulting amino acids or dipeptides are then imported into the cytoplasm (Smirnova et al., 2012; Suzuki et al., 1986) (see Figure 3A for a schematic overviuew). Since cysteine has reductive power by itself, we wondered if imported cysteine or cysteine-containing dipeptides caused the observed Grx1-roGFP2 reduction in cells lacking both endogenous glutathione and the Gsi transporter. To address this, we additionally deleted the gene coding for Ggt in these cells and analyzed reduction of Grx1-roGFP2 by exogenous GSH in the resulting Δ*gshA* Δ*gsiC* Δ*ggt* triple deletion mutant. External GSH still reduced Grx1-roGFP2 after addition, with a velocity virtually identical to the Δ*gshA* Δ*gsiC* strain or the Δ*gshA* single mutant (Figure 3B). This suggests that cysteine or cysteine-containing dipeptides derived from glutathione hydrolysis are not responsible for the reduction of Grx1-roGFP2 under these conditions.

**Figure 3.**
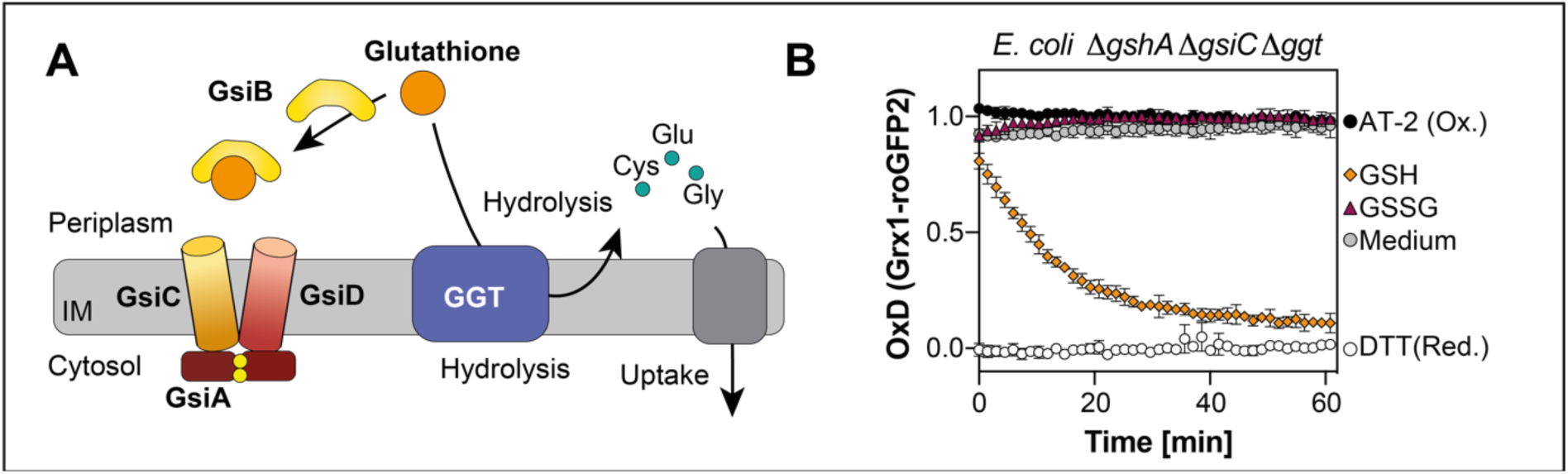
Glutathione hydrolysis by Ggt and cysteine or cysteine-containing dipeptide import is not responsible for Grx1-roGFP2 reduction in glutathione-deficient cells lacking GsiC. **(A)** Schematic overview of glutathione hydrolysis by Ggt and import of the hydrolysis products. **(B)** Grx1-roGFP2 oxidation state expressed in *E. coli* Δ*gshA* Δ*gsiC* Δ*ggt*. Probe was expressed and probe oxidation was recorded as described in Figure 1. Oxidized (AT-2) and reduced (DTT) cells were used for calculation of the probe’s oxidation degree (OxD). Plotted values represent the mean of three individual experiments, error bars represent the standard deviation. All experiments were performed as technical triplicates.

### The oligopeptide transporter OPP imports reduced glutathione into the cytosol of E. coli

Based on our findings that the glutathione redox homeostasis can be fully restored by the addition of endogenous reduced glutathione to a Δ*gshA* mutant lacking the transporter Gsi, essentially with the same velocity as in a mutant lacking only GshA, we hypothesized the presence of a yet unidentified glutathione transporter specific for reduced glutathione.

We thus attempted to identify ABC transporters through a similarity search. Using GsiB as a template, a BLAST search against the *E. coli* proteome identified several candidates for periplasmic binding proteins of ABC transporters. Among them were the periplasmic binding proteins of more or less well-characterized ABC transporters, such as DppA with 49% similarity, known to import di-and tripeptides (Kuenzl et al., 2018; Smith et al., 1999), SapA with 43% similarity, known for the transport of antimicrobial peptides in *Salmonella typhimurium* (Parra-Lopez et al., 1993) and *Haemophilus influenza* (Shelton et al., 2011) and the export of putrescine in *E. coli* (Sugiyama et al., 2016), OppA (43% similarity), part of an oligopeptide transporter (Klepsch et al., 2011), and YejA (36% similarity), a putative oligopeptide transporter, which is known to confer resistance to the peptide antibiotic microcin C (Novikova et al., 2007; Vondenhoff et al., 2011) (see Figure 4A for a schematic overview). In order to test, if one of these ABC transporters is needed to import reduced glutathione, we deleted genes coding for one of the permease domains (OppC, DppC and SapC) or the periplasmic binding protein in case of Yej (YejA) in cells lacking glutathione biosynthesis and the GsiC permease. We then expressed Grx1-roGFP2 in the cytosol of the resulting mutants and analyzed probe reduction by external GSH. Only the lack of OppC and GsiC in GSH-deficient cells (Δ*gshA* Δ*gsiC* Δ*oppC*) completely inhibited GSH-depended reduction of Grx1-roGFP2, whereas the other triple mutants were unaffected (Figure 4B-E). These experiments suggest that Opp is a highly efficient transporter for reduced glutathione. However, as our previous experiments with the Δ*gshA* Δ*gsiC* double mutant have shown, Opp cannot transport oxidized GSSG (Figure 2B-D).

**Figure 4.**
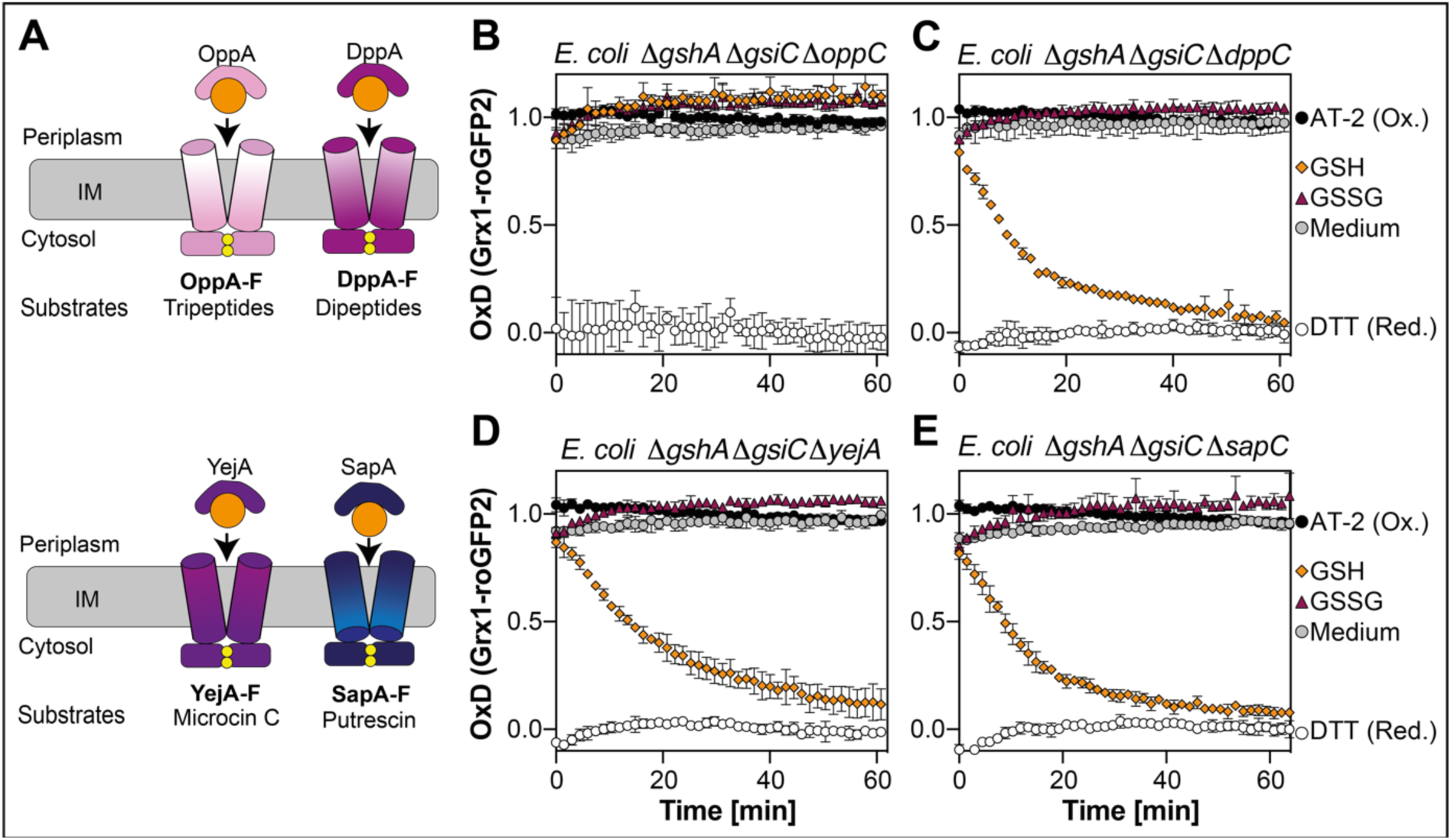
The OppA-F transporter imports reduced glutathione. **(A)** Schematic overview of known peptide transporters in *E. coli*. Oxidation state of Grx1-roGFP2 expressed in *E. coli* triple deletion strains lacking GSH biosynthesis, GsiC and one domain of putative GSH transporters: Δ*gshA* Δ*gsiC* Δ*oppC* **(B)**, Δ*gshA* Δ*gsiC* Δ*dppC* **(C)**, Δ*gshA* Δ*gsiC* Δ*yejA* **(D)** or Δ*gshA* Δ*gsiC* Δ*sapC* **(E)**. The probe was expressed and oxidation state measured as described in Figure 1. Oxidized (AT-2-) and reduced (DTT-treated) cells were used for calculation of the probe’s oxidation degree (OxD). Plotted values represent the mean of three individual experiments, error bars represent the standard deviation. All experiments were performed as technical triplicates.

### Lack of Opp periplasmic binding protein OppA or permease OppC results in decrease of GSH-dependent reduction of Grx1-roGFP2 in GSH-deficient cells

Thus far, our data suggested to us that Gsi is the only permease capable of importing oxidized glutathione into the cytoplasm and that Opp is a highly efficient permease for reduced glutathione. The fact that there was no discernible difference in the velocity in Grx1-roGFP2 reduction by exogenous reduced glutathione in a Δ*gshA* mutant lacking or containing Gsi suggested to us that Opp is the main transporter for reduced glutathione in *E. coli*.

To test our hypothesis, we deleted genes coding for the periplasmic binding protein OppA or the permease OppC in cells lacking glutathione biosynthesis, creating double mutants Δ*gshA* Δ*oppA* and Δ*gshA* Δ*oppC* (see Figure 5A for a schematic overview). Subsequently, we analyzed reduction of Grx1-roGFP2 in these cells by exogenous oxidized or reduced glutathione. The observed reduction of Grx1-roGFP2 in both mutant cells by oxidized glutathione was comparable to the Δ*gshA* single mutant, concurrent with our finding that Gsi is the sole transporter for GSSG. However, reduction by external GSH in both strains was noticeably slower and started even later than probe reduction by GSSG (Figure 5B, C).

**Figure 5.**
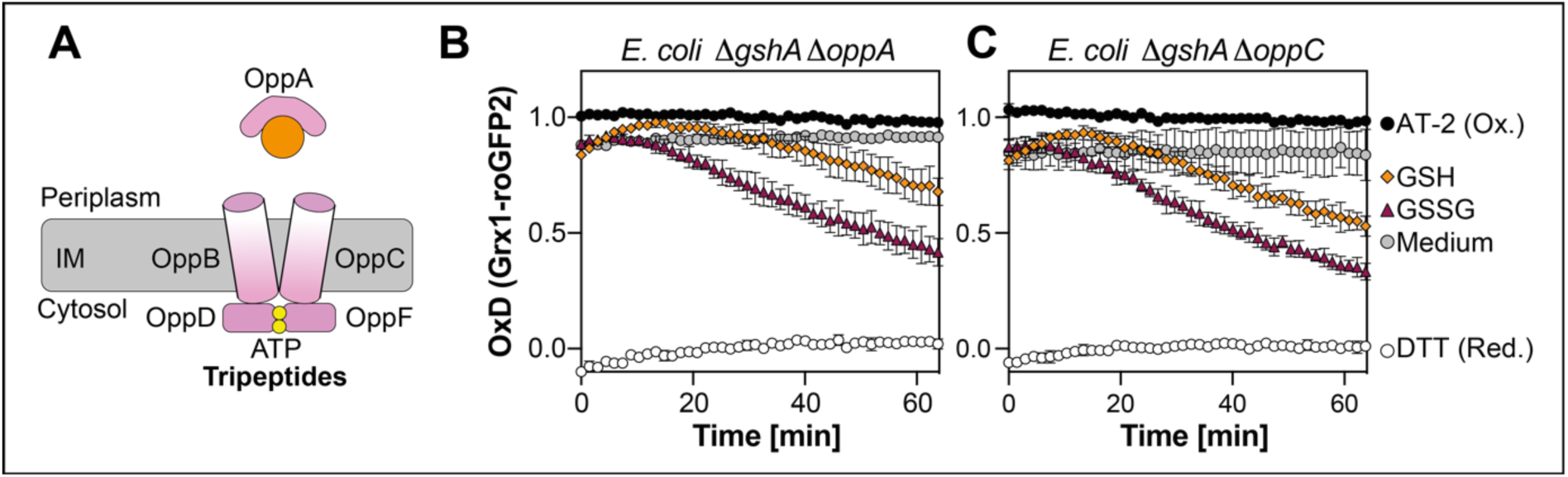
Import of reduced glutathione is significantly slower in GSH-deficient cells lacking OppA or OppC. **(A)** Schematic of *E. coli* OppA-F ABC transporter. Oxidation state of Grx1-roGFP2 expressed in *E. coli* Δ*gshA* Δ*oppA* **(B)** or Δ*gshA* Δ*oppC* **(C)**. Grx1-roGFP2 expression and analysis of probe oxidation was performed as described in Figure 1. Oxidized (AT-2-) and reduced (DTT-treated) cells were the controls for calculation of the probe’s oxidation degree (OxD). Plotted values represent the mean of three individual experiments, error bars represent the standard deviation. All experiments were performed in triplicate measurements.

This suggested to us that Opp is indeed the major transporter for reduced glutathione in *E. coli*. Since reduced glutathione has a half-life of around 9 hours at pH 7.5 in the absence of metals (Stevens et al., 1983) and multiple times faster in the presence of certain metals (Voegtlin et al., 1931), we wanted to exclude that GSH oxidizes over time in our assay, resulting in the formation of GSSG. This accumulating GSSG could then be taken up by Gsi, even if Gsi was specific only for GSSG. To prevent metal-dependent GSH oxidation, we added 5 mM EDTA or GSH and EDTA to cell suspensions of Δ*gshA* and Δ*gshA* Δ*oppC* (see Figure 6A for a schematic overview). However, we did not observe a significant effect of EDTA on GSH-dependent reduction of the probe expressed in these cells (Figure 6B, C). We also added glutathione reductase Gor (1 µM) and its co-substrate NADPH+H^+^ (250 µM), to ensure the retention of the reduced state of the exogenous glutathione (see Figure 6D for a schematic overview). There was no significant retardation of probe reduction in either Δ*gshA* or Δ*gshA* Δ*oppC* cells, at least when compared to the addition of Gor without NADPH (Figure 6E, F). Unless the oxidizing environment of the periplasm induces the formation of GSSG our data suggests that Gsi is capable of transporting reduced glutathione, albeit significantly less efficient than GSSG.

**Figure 6.**
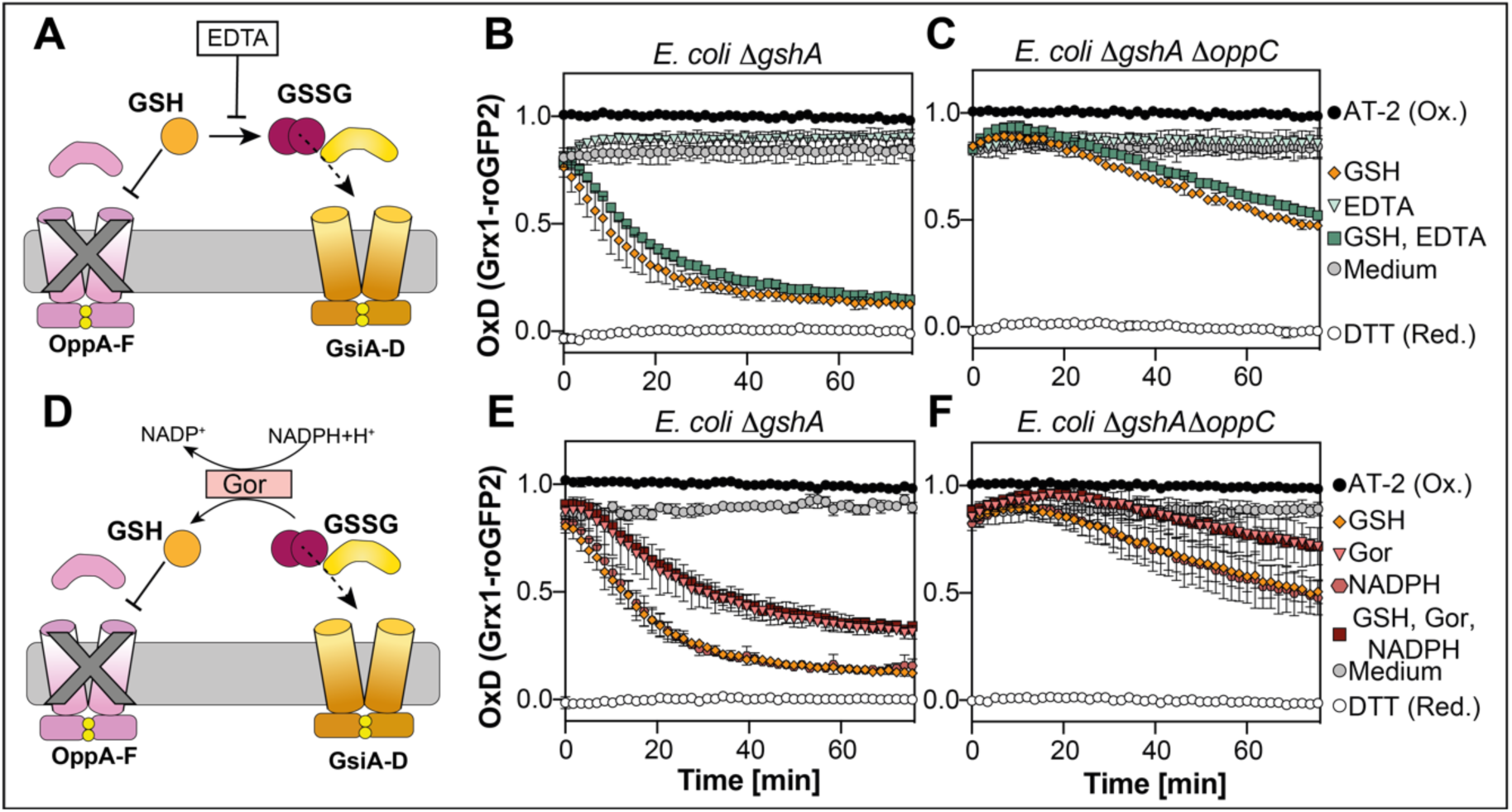
External addition of EDTA or Gor and NADPH does not prevent Grx1-roGFP2 reduction by GSH in GSH-deficient cells lacking OppC. Schematic overview of EDTA addition to prevent GSH oxidation **(A)** or Gor-based reduction of GSH **(D)**. Grx1-roGFP2 reduction in the presence of 5 mM EDTA in Δ*gshA* (B) or Δ*gshA* Δ*oppC* (E) or 1 µM Gor and 250 µM NADPH+H^+^ in Δ*gshA* (C) or Δ*gshA* Δ*oppC* (F). Cells were cultivated and Grx1-roGFP2 expressed as described in Figure 1. Grx1-roGFP2 fluorescence was recorded as described in Figure 1 and AT-2-(oxidized) and DTT-treated (reduced) cells served as controls for OxD calculation. Plotted values represent the mean of three individual experiments, error bars represent the standard deviation. All experiments were performed as technical triplicates.

### The periplasmic binding proteins of the Opp and Gsi transporters do not cross react with the respective permease domains

Studies with *Haemophilus influenza* have shown, that HbpA, a designated periplasmic binding protein transfers glutathione to the Dpp-like dipeptide ABC transporter, while DppA, the binding protein of Dpp does not bind glutathione (Vergauwen et al., 2010). In order to investigate if the substrate binding protein of Gsi (GsiB) transfers GSH to Opp or *vice versa* (OppA to Gsi), we generated GSH-deficient cells lacking either OppA and GsiC or GsiB and OppC and expressed Grx1-roGFP2 in the resulting Δ*gshA* Δ*gsiB* Δ*oppC* and Δ*gshA* Δ*oppA* Δ*gsiC* triple mutants (see Figure 7 A for a schematic overview). Cytoplasmic Grx1-roGFP2 was not reduced upon the addition of either reduced or oxidized glutathione in both Δ*gshA* Δ*gsiB* Δ*oppC* and Δ*gshA* Δ*oppA* Δ*gsiC* strains. This shows that there is no cross-talk between the periplasmic binding proteins and the opposite permeases and also supports our hypothesis that Gsi and Opp are the only transporters capable of importing exogenous glutathione into the cytoplasm (Figure 7B, C).

**Figure 7.**
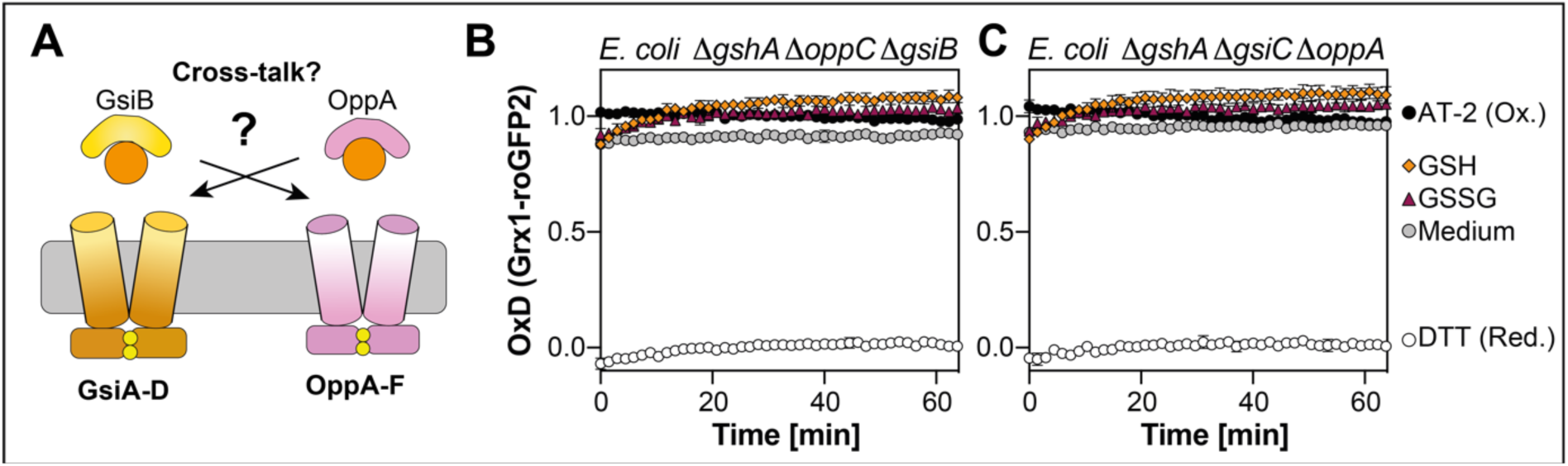
Import of reduced or oxidized glutathione does not depend on crosstalk of periplasmic binding proteins with the permeases. **(A)** Schematic overview of postulated crosstalk between periplasmic binding proteins with permeases. Oxidation state of Grx1-roGFP2 expressed in *E. coli* Δ*gshA* Δ*gsiB* Δ*oppC* **(B)** or Δ*gshA* Δ*oppA* Δ*gsiC* **(C)**. Cells were cultivated and Grx1-roGFP2 oxidation was determined as described in Figure 1. For calculation of the probe’s oxidation degree (OxD), oxidized (AT-2-) and reduced (DTT-treated) cells were used. Plotted values represent the mean of three individual experiments, error bars represent the standard deviation. Individual experiments were performed as technical triplicates.

## Discussion

The tripeptide glutathione is a major biological antioxidant and essential in a variety of organisms. GSH forms GSSG upon oxidation, which in turn is reduced back by glutathione reductase in an NADPH-dependent mechanism (Couto et al., 2016; Greer and Perham, 1986; Williams and Arscott, 1971). Glutaredoxins (Grxs) can reduce disulfides formed in cytosolic proteins under oxidative and nitrosative stress, and GSH recycles oxidized Grxs (Holmgren, 1979, 1976; Lillig et al., 2008; Vašková et al., 2023), making glutathione one of the most important thiol-disulfide redox co-factors. Here, we used the Grx1-roGFP2 redox probe in the cytosol of cells lacking the first enzyme of glutathione biosynthesis to dissect glutathione import in the model organism *Escherichia coli*. The fact that this probe was highly oxidized in the cytosol of these cells made it a valuable tool to monitor glutathione import: addition of both, oxidized or reduced glutathione recovered Grx1-roGFP2 redox state. Reduction by GSSG was significantly slower, but not absent, showing that GSH-deficient cells can use glutathione reductase to restore their highly reductive cellular glutathione redox state from the oxidant GSSG. These findings underline the importance of the glutathione reductase for maintenance of the GSH/GSSG homeostasis, as recently reviewed in (Couto et al., 2016).

Albeit synthesized in the cytosol, GSH is actively secreted and imported by exponentially growing bacteria. Recent findings suggested that the ABC transporter Gsi is the only transporter specific for intact glutathione across the inner membrane from the periplasm into the cytosol in *Escherichia coli*, since uptake assays with GshA, Gsi and Ggt deficient cells showed only trace amounts of intracellular glutathione in a study by (Suzuki et al., 2005) and even no glutathione uptake was observed in a study by (Wang et al., 2018) in these cells. Here, we provide evidence that this view of glutathione transport in *E. coli* needs to be revisited. Our data, based on reactions of the genetically encoded redox probe Grx1-roGFP2 suggests that there are two, and only two systems capable of glutathione import into the cytoplasm of *E. coli*. One is indeed Gsi and our data strongly suggests that its preferred substrate is oxidized and not reduced glutathione. The bulk of reduced glutathione, however, is transported by the Opp transporter. Both transporters depend on their periplasmic binding protein for glutathione transport and there is no crosstalk between the transport systems.

We excluded that hydrolysis products of glutathione were responsible for the observed reduction, since Grx1-roGFP2 in Δ*gshA* Δ*gsiC* mutant strains lacking Ggt (Smirnova et al., 2012; Suzuki et al., 1986), could still be reduced by glutathione. In line with that, in a previous study investigating extracellular glutathione depletion in cells lacking *gshA*, *ggt*, and *gsiAB,* glutathione accumulation in the cells was not completely absent, however drastically reduced (Suzuki et al., 2005). This study relied on radioactively labeled glutathione, and to our knowledge, did not control for glutathione oxidation, which could explain the reduced uptake of glutathione in those strains. A recent study aimed to characterize the periplasmic binding protein GsiB and suggested that GsiB binds both, reduced and oxidized glutathione, however the authors did not provide any data on binding affinity or kinetics (Wang et al., 2018). Our data suggests that Gsi might transport reduced glutathione, but at a much lower rate than its oxidized form and certainly not as effective as Opp. The fact that (Suzuki et al., 2005) and (Wang et al., 2018) did observe little or no import of glutathione could be due to oxidation of the exogenous glutathione.

Although we showed that neither glutathione reductase, nor EDTA prevented Grx1-roGFP2 reduction by exogenous reduced glutathione in a Δ*gshA* Δ*oppC* strain, we can, however, not exclude that Gsi might be specific for oxidized glutathione. GSH in solution forms GSSG over time (Stevens et al., 1983) and our efforts with the addition of EDTA or glutathione reductase might have fully inhibited this oxidation, but before entering the cytosol, GSH also needs to pass the periplasm, a highly oxidative environment (Knoke et al., 2023; Manta et al., 2019). In this compartment GSH might get oxidized, especially, when direct GSH import is abolished, and the resulting GSSG could then be transported by Gsi.

The presence of two separate transporters for reduced and oxidized glutathione enables cells to regulate glutathione uptake. This might be important when bacteria encounter elevated extracellular oxidant concentrations, for example during host colonization, since immune cells produce reactive oxygen species to kill invading pathogens (Degrossoli et al., 2018; Xie et al., 2019). In line with this, OppA is known as virulence factor in many Gram-negative and Gram-positive bacteria (Tam and Saier, 1993; Zheng et al., 2018), such as *Streptococcus suis*. In pathogenic mycobacteria, a recently identified OppA-like glutathione binding protein has been shown to modulate the immune response of infected macrophages (Dasgupta et al., 2010). Additionally, Opp mutants of *Mycobacterium bovis* (Green et al., 2000), *Corynebacterium pseudotuberculosis* (Moraes et al., 2014) or *Vibrio alginolyticus* (Liu et al., 2017) were more susceptible towards glutathione and showed reduced virulence. This highlights the importance of the role of glutathione transport and its regulation in the host-microbe interplay. Concomitantly, previous and our data suggest that the promiscuous Opp oligopeptide transporter is an ubiquitous GSH import system, however, whether or not Opp in other organisms imports oxidized glutathione needs to be elucidated. In any case, the presence of a dedicated transporter for oxidized glutathione, such as Gsi, could provide an additional benefit in the context of virulence: extracellular bodily fluids, such as blood plasma, typically contain high proportions of glutathione in its oxidized form.

## Conclusion

Based on our data, we propose a comprehensive model for glutathione import into the cytoplasm of *E. coli*: Gsi is the exclusive transporter for oxidized glutathione, and, if at all, transports reduced glutathione with a low efficiency, while Opp exclusively imports reduced glutathione (Figure 8).

**Figure 8.**
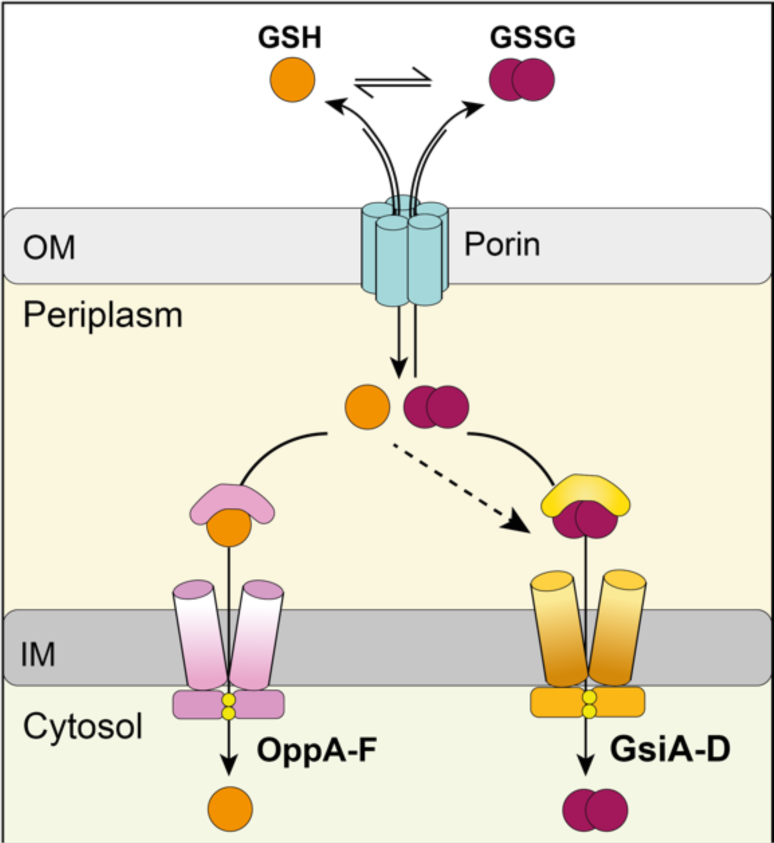
Model of the import of oxidized and reduced glutathione into *E. coli*’s cytoplasm. Our data suggests that GSSG is exclusively transported by GSI, and that OPP imports the bulk of extracytoplasmic GSH. OM, outer membrane; IM, inner membrane.

## Acknowledgements

We thank Franz Narberhaus (Ruhr University Bochum) for kindly providing the 709-FLPe plasmid.

## Author contributions

LRK and LIL designed the study. LRK and MM planned the experiments. LRK and MM constructed the double and triple deletion strains. MM performed the Grx1-roGFP2 measurements and LRK and MM evaluated the generated data. LRK and LK performed initial experiments. LRK and LIL carried out the data interpretation. LRK wrote the manuscript. LIL edited the manuscript. All authors consulted on the manuscript and approved the final version.

## Data availability statement

The data supporting the findings of this study is presented within the article. Strains and plasmids constructed for this study are available upon request.

